# IsoMaTrix: a framework to visualize the isoclines of matrix games and quantify uncertainty in structured populations

**DOI:** 10.1101/2020.06.24.170183

**Authors:** Jeffrey West, Yongqian Ma, Artem Kaznatcheev, Alexander R. A. Anderson

## Abstract

**Summary:** Evolutionary game theory describes frequency-dependent selection for fixed, heritable strategies in a population of competing individuals using a payoff matrix, typically described using well-mixed assumptions (replicator dynamics). IsoMaTrix is an open-source package which computes the isoclines (lines of zero growth) of matrix games, and facilitates direct comparison of well-mixed dynamics to structured populations in two or three dimensions. IsoMaTrix is coupled with a Hybrid Automata Library module to simulate structured matrix games on-lattice. IsoMaTrix can also compute fixed points, phase flow, trajectories, velocities (and subvelocities), delineated “region plots” of positive/negative strategy velocity, and uncertainty quantification for stochastic effects in structured matrix games. We describe a result obtained via IsoMaTrix’s spatial games functionality, which shows that the timing of competitive release in a cancer model (under continuous treatment) critically depends on the initial spatial configuration of the tumor.

**Availability and implementation:** The code is available at: https://github.com/mathonco/isomatrix.

## Introduction

Interactions between competing individuals which result in some benefit or cost can broadly be described and analyzed using a mathematical framework called game theory. This framework aims to mathematically determine the optimal strategy to employ when in competition with an adversary^1^. The components of a game are: 1) the strategies, 2) the players, and 3) the payoffs of each strategy. This classical definition of game theory can be extended to model evolution by natural selection, known as evolutionary game theory (EGT)^2, 3^. Typically, EGT describes changes in the prevalence of strategies due to frequency-dependent selection in a population of competing players that adhere to fixed strategies (but see ref. 4 for alternative experiment-focused interpretation of the basic terms of EGT). Competition is governed by a “payoff matrix,” defining the Darwinian fitness of an individual based upon interactions with other individuals within the population.

EGT is increasingly used to model cancer as an evolutionary process^5, 6^, and viewed as one of the central paths forward in the roadmap of mathematical oncology^7^. EGT models have shown success in modeling glioma progression^8^, tumor-stroma interactions^9^, growth factor production as a public good^10, 11^, effects of tissue edges on cancer cell motility^12^, tumor growth^13^, vascularization and tumor acidity^14^, metastatic prostate cancer and the bone-remodelling cycle^15^, competitive release^16, 17^, and optimal cancer treatment^18, 19^. Recently, the payoff matrices that specify evolutionary games have even been directly measured *in vitro*^20^.

To help mathematical oncologists along the path of EGT modeling, we develop a software package to systematically analyze three-strategy matrix games. The package allows for comparison between analysis of the non-spatial, well-mixed assumption (i.e. replicator equation) to spatially-explicit formulations of matrix games in two- or three-dimensions. The package places a special focus on boundaries between the positive and negative growth regions of each strategy, known as isoclines. Thus, the name of this package, IsoMaTrix, is a blend of “isocline” and “matrix” games, to describe this key functionality.

### Distinguishing features of Isomatrix

Several groups have released similar packages in Mathematica that compute evolutionary game dynamics of three strategy games (Dynamo^21, 22^, EvoDyn-3s^23^). More recently, EGTplot (Python) allows for static or animated images of dynamics^24^. An extension to model multiplayer games with collective interactions (public goods games) was designed in Mathematica (DeFinetti)^25^.

The foremost distinguishing feature of the IsoMaTrix package is the extension of explicit spatial structure, including functions to quantify and visualize uncertainty due to stochastic effects. This extension, facilitated by a Hybrid Automata Library (HAL^26^; java language) module, allows for easy comparison between non-spatial (replicator dynamics) and spatial (cellular automata) model configurations. Other distinguishing features include computation of isoclines, novel region plots, simultaneous visualization of multiple games, and automatic computation of basins of attraction.

## Methods

### Replicator dynamics

Frequency-dependent selection dynamics in IsoMaTrix are governed by the replicator equation:

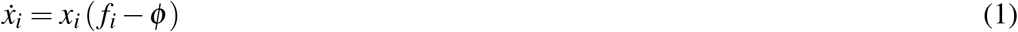

where *x*_*i*_ is the fraction of each strategy, *f*_*i*_ = ∑ _*j*_ *a*_*i j*_*x* _*j*_ is the fitness of each strategy, and the average fitness is *ϕ* = ∑ _*j*_ *f* _*j*_*x* _*j*_ (with *i, j ∈* [1, 2, 3]). The fraction of each strategy grows or decays with exponential rate proportional to its relative fitness: i.e. the fitness difference above/below the average fitness of the population, *ϕ*. We consider three strategy games with payoff matrix, *A* such that [*A*]_*i j*_ = *a*_*i j*_.

The IsoMaTrix package has functions which display 1) fixedpoints, 2) isoclines, 3) phase flow, 4) velocities, 5) trajectories, 6) regions of positive/negative strategy fitness and 7) basins of attraction. Each function is independently called, enabling chaining to facilitate the desired visualization of dynamics. Figure 1 shows a representative example of visualizations possible in IsoMaTrix.

**Figure 1.**
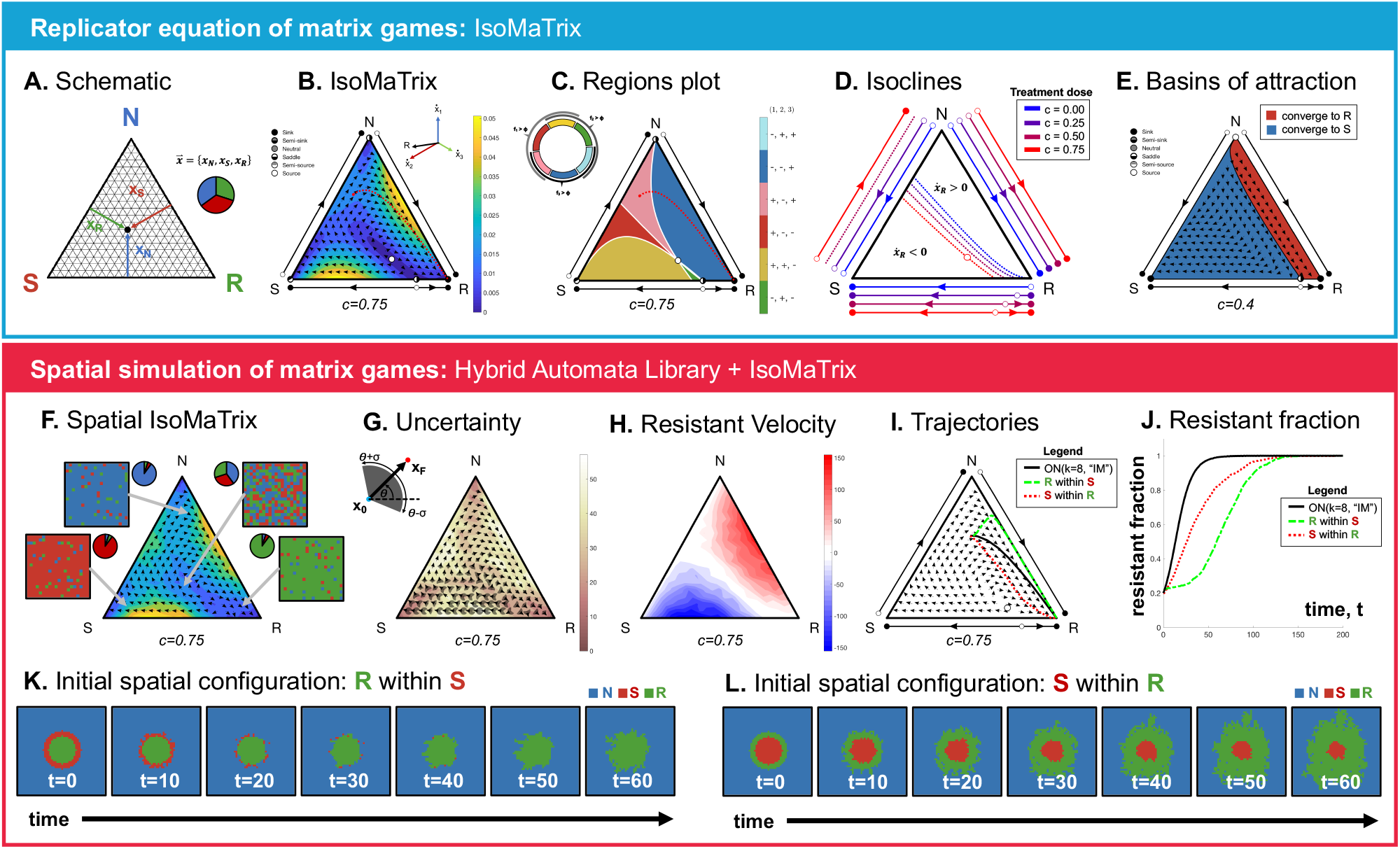
IsoMaTrix. Top section: well-mixed dynamics. (A) Schematic of simplex describing competition between Normal (N), Sensitive (S), and Resistant (R) cell types. (B) IsoMaTrix diagram for payoff matrix in eqn. 2. (C) Region plot, with regions delineated by positive/negative strategy velocity, 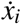. Example trajectory starting from 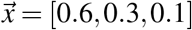, where the tumor initially decays (negative sensitive velocity: 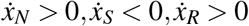; pink) but quickly relapses with saturation of the resistant type (positive resistant velocity: 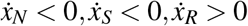; blue). (D) Isoclines for resisant strategy 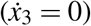 for varied dose, *c* ∈ [0, 0.25, 0.5, 0.75] (blue to red). Bottom section: spatial dynamics. (E) Phase flow shown for *c* = 0.25, with simulated spatial structure using the Ohtsuki-Nowak transform (*k* = 4, Death-Birth updating). Color-coded by basins of attraction. (F) Schematic of IsoMaTrix diagrams computation in structured populations. *M* stochastic realizations are simulated for each initial proportion (i.e. 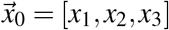) within a mesh covering the triangle: four examples shown inset. Phase flow is estimated by calculating the resultant vector between 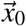 and the final proportion, 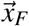, after user-specified *n* number of time steps. (G) Uncertainty: standard deviation, *σ*, of the magnitude (background-color) and of direction (gray arc on each phase-flow arrow). (H) Subvelocity resistant cells. (I,J) The trajectories through space (I) and time (J) for configurations shown in (K,L) compared to well-mixed (black). (K,L) Two initial spatial configurations over time. Resistant cells trapped within the core (top) delays the emergence of resistant cells under treatment.

### Importance of spatial structure

While replicator dynamics has proven quite useful, the dynamics of spatially structured populations can vary dramatically^27, 28^. In some cases, it is possible to create transforms of the replicator dynamics due to the specific spatial structures. The effect of space is then equivalent to an invertible transformation of the entries in the payoff matrix^12, 29^–32. IsoMaTrix provides an implementation of one such analytic transform^29^ to get modelers started with first-order deviations from well-mixed dynamics due to spatial structure.

IsoMaTrix uses the on-lattice cellular automata framework HAL^26^ to perform simulation-based analysis of spatial structure. IsoMaTrix uses ‘imitation updating’ whereby a randomly chosen focal cell updates its strategy to imitate one of its own neighbors in proportion to fitness^29, 33^. Either deterministic (maximum fitness neighbor strategy wins) or stochastic (weighted by fitness of all neighbors) updating can be chosen^34, 35^.

In summary, IsoMaTrix allows for quick analysis of well-mixed matrix games in an accessible language (MATLAB), as well as detailed comparison of the effect of spatial structure on dynamics.

### Example

To illustrate the utility of IsoMaTrix, figure 1 uses a simplified version of the payoff matrix from ref. 16, used to model competitive release:

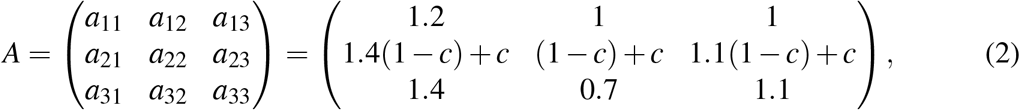

where the rows and columns describe competition between Normal (N; first row and column), Sensitive (S; second), and Resistant (R; third) cells within the tumor bed. Competition depends on drug concentration, *c*.

The top section of figure 1 shows an example IsoMaTrix output for non-spatial EGT matrix games using the replicator equation. Dynamics are displayed on a triangular plot (figure 1A) where each corner represents a tumor with 100% of the given strategy. Figure 1B shows phase flow under treatment (black arrows), with background colored by velocity magnitude. Fixed points for each pair-wise strategy interaction (N-S, S-R, N-R) are conveniently offset on each edge with arrows indicating phase flow (black lines; solid circles are stable while open circles are unstable). In figure 1C, the aforementioned ‘regions’ plot delineates the ternary plot into color-coded regions according to if each strategy (*i ∈*{1, 2, 3}) is selected for (*f*_*i*_ > *ϕ*) or against (*f*_*i*_ < *ϕ*). Knowledge of the resistant isocline, shown changing with dose strength in figure 1D, facilitates control of the tumor dynamics by allowing treatment to be discontinued well before resistant regrowth^13, 16, 18^. Importantly, IsoMaTrix allows for multiple games (in this case, multiple values of dose) to be easily displayed on the same ternary diagram.

The bottom section of figure 1 shows example IsoMaTrix output for spatial simulations governed by the same payoff matrix, A. Figure 1F shows a schematic detailing how IsoMaTrix diagrams are produced for structured populations. *M* stochastic realizations are simulated for each initial proportion (i.e. 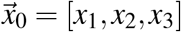) within a mesh covering the triangle. The phase flow is estimated by calculating the resultant vector between and the initial and final proportion after *n* time steps. Given the stochastic nature of spatial simulations, uncertainty can be calculated (figure 1G).

IsoMaTrix facilitates comparison of precise spatial configurations. Figures 1I-L compare the well-mixed (black line) dynamics to two spatial configurations: resistant cells trapped inside sensitive cells (K) and vice versa (L). The tumor with resistant cells trapped within the core delays the emergence of resistant cells (figure 1J) under treatment.

### IsoMaTrix Manual

#### 1 IsoMaTrix (MATLAB)

Each of the following subsections corresponds to a function declaration in the IsoMaTrix package. Unless otherwise noted, the following payoff matrix is used to describe competition between strategy 1 (first row/column), 2 (second row/column), and 3 (third row/column):

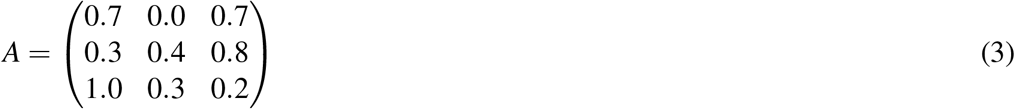

Colors are specified consistent with MATLAB conventions: a 1×3 vector, [*R, G, B*], where each vector element is [0, 1]. For example, the following colors are used in subsequent code:

~~~
black=[0,0,0];
red=[1,0,0];
green=[0,1,0];
blue=[0,0,1];
~~~

##### 1.1 isomatrix(A)

Isomatrix is the base function in the package:

~~~
isomatrix(A);
~~~

This function reads in a payoff matrix, *A*, and automatically generates a series of 5 figures, corresponding to each panel in figure 2:

**Figure 2.**
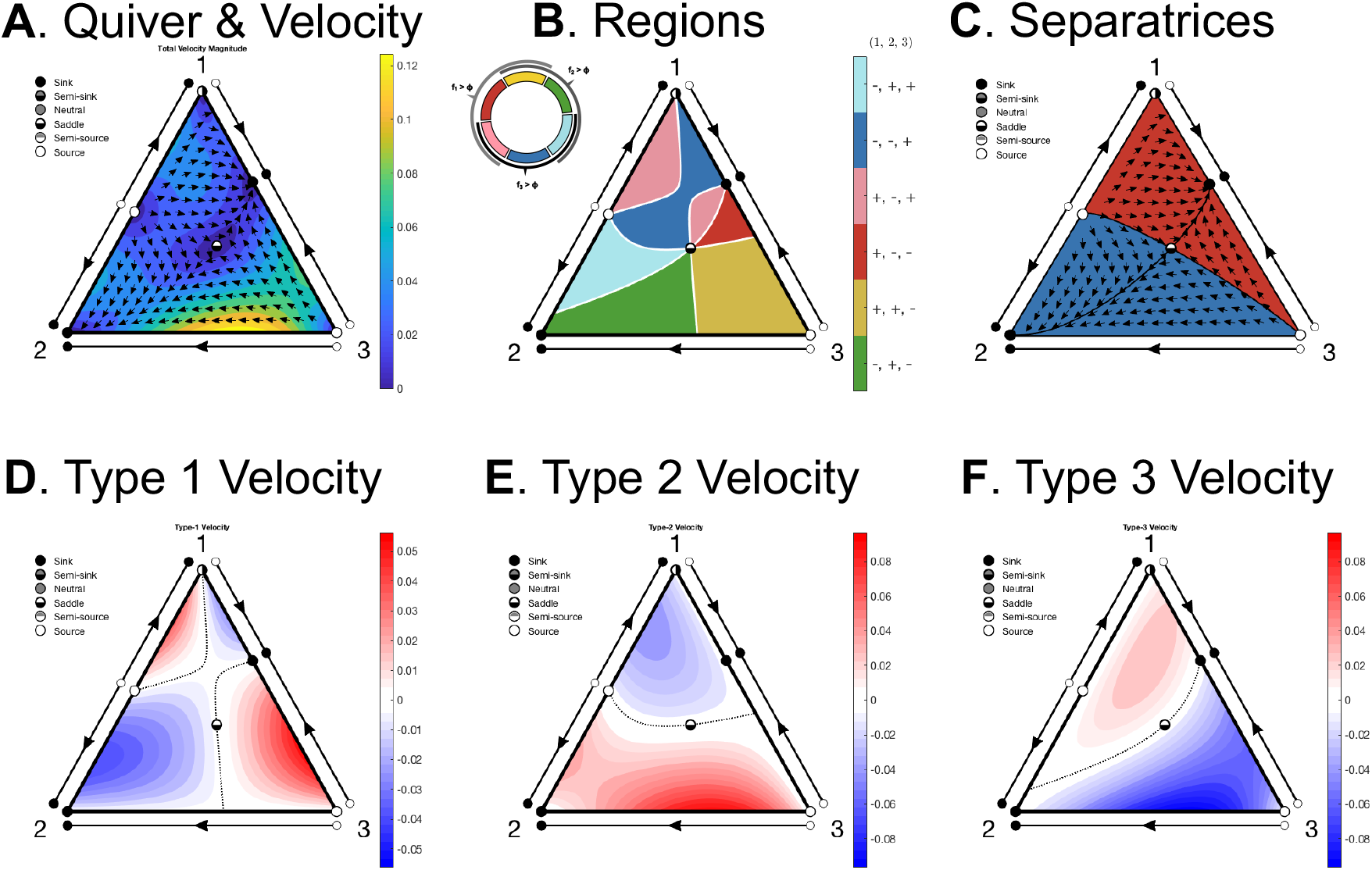
IsoMaTrix’s base function. The function, “isomatrix” produces five diagrams derived from the input payoff matrix, *A*. (A) A quiver plot (black arrows) shows the phase flow, where the background color indicates the magnitude of the velocity vector. (B) A “isomatrix_region” plot divides the diagram into regions of positive or negative growth of each strategy (delineated by the strategy isoclines in white), with the signs indicated by colorbar. For example green indicates the region where 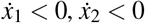, and 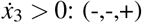. (C,D,E) Strategy velocity magnitude is color-coded by positive (red) or negative (blue) growth of each strategy, with nullcline shown in black. Pairwise fixed points are also drawn on each edge.

**A** quiver & fixed points & total velocity

**B** fixed points & “regions” plot

**C** fixed points & subvelocity for strategy 1

**D** fixed points & subvelocity for strategy 2

**E** fixed points & subvelocity for strategy 3

An example output for payoff matrix *A* (eqn. 3) is shown in figure 2. Each of these functions (quiver plots, fixed points, velocity, and subvelocity) can be plotted separately, and are explained in the following sections. If the user desires to label the simplex corners, ‘Labels’ is an optional name-value argument:

~~~
isomatrix(A,’Labels’,{‘1’,’2’,’3’});
~~~

##### 1.2 isomatrix_fixedpoint(A,index)

The fixed point function draws the pairwise interaction lines on each simplex edge. Solid circles represent stable fixed points while open circles represent unstable fixed points. If an internal fixed point exists, it is drawn with a square. The function takes as arguments a 3×3 payoff matrix, *A* and (optionally) an index (integer; default value of 1) indicating the distance from the simplex edge.

~~~
isomatrix_fixedpoint(A);
~~~

If an index is specified, only the edge fixed points are displayed (offset), along with any internal fixed points (s.t. `min`(*x*_*i*_) > 0). If an index is not specified, corner and edge fixed points are displayed as (semi)sink, (semi)source, saddle, or neutral (see e.g. figure 2A).

Other optional name-value arguments are ‘Color’ and ‘Labels.’ Shown in figure 3A are the fixed points calculated for *A* in eqn. 3 (black), *A*^*T*^ (red), and 1 − *A* (blue). This plots fixed point diagrams on each edge, offset by the index value. This can be accomplished with the following code:

~~~
isomatrix_fixedpoint(A,1,’Color’,black,’Labels’,{‘1’,’2’,’3’});
isomatrix_fixedpoint(A’,2,’Color’,red);
isomatrix_fixedpoint(1-A,3,’Color’,blue);
~~~

**Figure 3.**
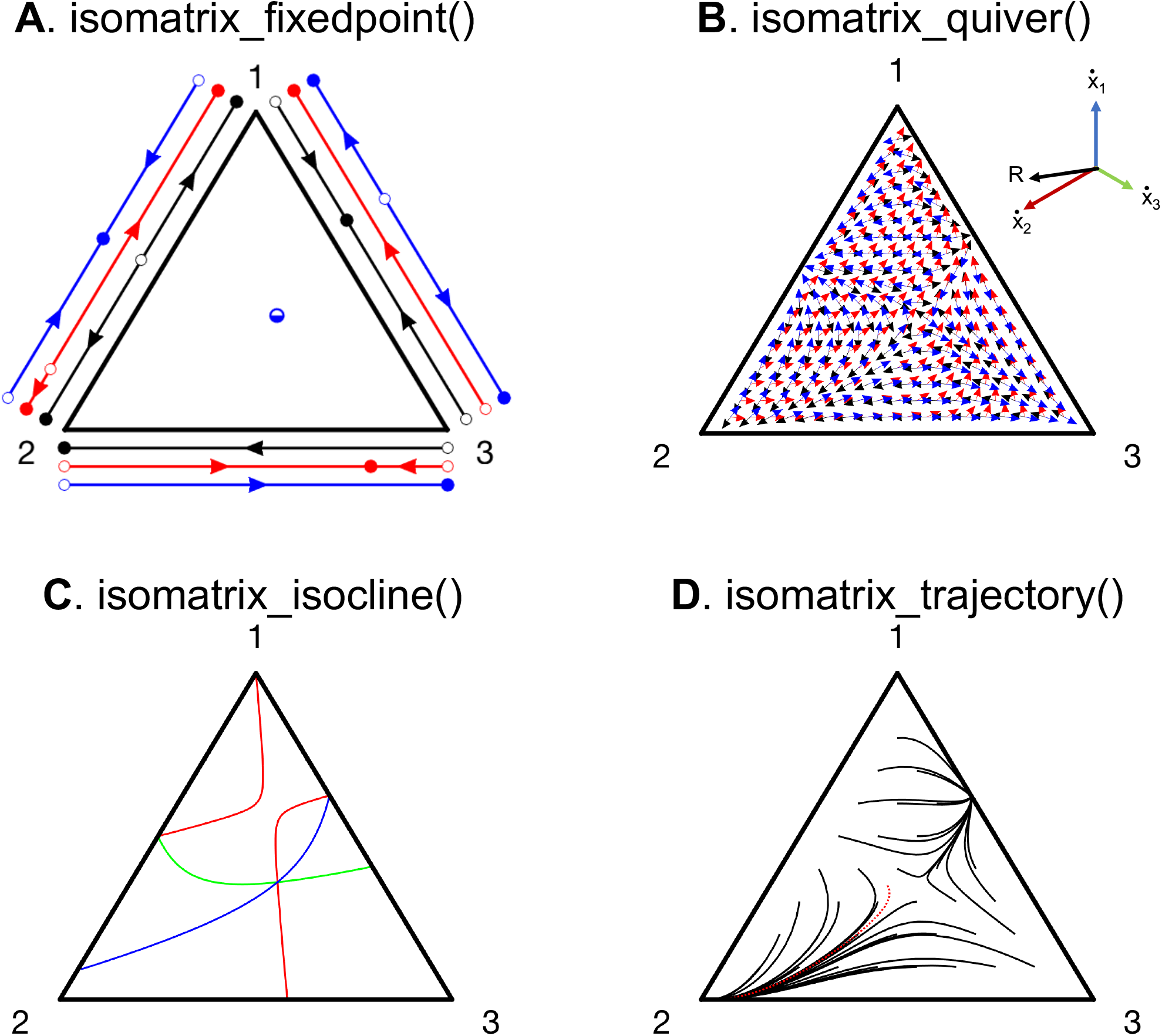
Additional IsoMaTrix Functions. (A) Output for “isomatrix_fixedpoint” function displays pairwise fixed points on each edge (closed circle for stable; open circle for unstable). (B) Output for “isomatrix_trajectory” function displays trajectories of matrix games. (C) Output for “isomatrix_isocline” displays the isocline for each strategy. (D) Output for “isomatrix_quiver” displays velocity vector field for matrix games.

##### 1.3 isomatrix_quiver(A)

The quiver function draws the phase flow, with evenly spaced arrows indicating the instantaneous velocity direction at each point (fig. 3B, inset). The function takes as an argument a 3×3 payoff matrix *A*.

~~~
isomatrix_quiver(A);
~~~

Other optional name-value arguments are ‘Color’ and ‘Labels.’ A schematic of the resultant arrow is shown inset in figure 3B.

~~~
isomatrix_quiver(A,’Color’,black,’Labels’,{‘1’,’2’,’3’});
isomatrix_quiver(A’,’Color’,red);
isomatrix_quiver(1-A,’Color’,blue);
~~~

##### 1.4 isomatrix_isocline(A,id)

The isocline function draws the lines of zero growth (sometimes referred to as nullclines) for each strategy. Isoclines indicate the bounding line between positive and negative growth: 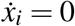. The function takes as arguments a 3×3 payoff matrix *A*, and a strategy id (between 1 and 3, inclusive). The id indicates the row/column of the strategy for which the isocline is calculated. If no id is specified, all three isoclines are shown in red, green, and blue.

~~~
isomatrix_isocline(A);
~~~

Other optional name-value arguments are ‘Color,’ ‘Labels,’ ‘LineWidth,’ and ‘LineStyle.’ The default setting is a red solid line of thickness 2. An example is shown in figure 3C for the following output:

~~~
isomatrix_isocline(A,1,’Color’, red, ‘Labels’,{‘1’,’2’,’3’} …
             ‘LineStyle’,’-’,’LineWidth’,2);
isomatrix_isocline(A,2,’Color’, green);
isomatrix_isocline(A,3,’Color’, blue);
~~~

##### 1.5 isomatrix_trajectory(A,x0,tF)

The trajectory functions plot one trajectory (or multiple trajectories) of a matrix game from a specified initial condition, 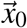. The initial condition is a n-by-3 array, where each row is a given initial condition. Replicator dynamics are simulated for *t*_*F*_ time-steps, and plotted on the IsoMaTrix diagram in user-specified color. If no initial condition is specified, trajectories are shown for initial conditions seeded uniformly across the domain (figure 3D, black lines) for 50 time-steps. An example single trajectory with user-specified initial condition is shown in red. Similar to isoclines, other optional name-value arguments are ‘Color,’ ‘Labels,’ ‘LineWidth,’ and ‘LineStyle.’

~~~
% evenly-distribution initial conditions (black):
isomatrix_trajectory(A);

% specified single initial condition (red):
tF=100;
x0=[0.3,0.3,0.4];
isomatrix_trajectory(A,x0,tF,’Color’,red, …
           ‘Labels’,{‘1’,’2’,’3’});
~~~

##### 1.6 isomatrix_velocity(A,id)

The purpose of this function is to color-code the background of the IsoMaTrix diagram according to the magnitude of velocity for the replicator dynamics. The function takes as arguments a 3×3 payoff matrix *A*, and a strategy id. The id should be an integer (i.e. 1, 2, or 3) to specify which strategy velocity to compute. Velocities are calculated directly from eqn. 1 with blue indicated negative velocities and red indicating positive. If no id is specified, the magnitude of the resultant velocity, ‖ *v* ‖, of all types is computed:

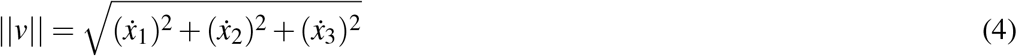

An example is shown in figure 5A. Examples of each strategy velocity is shown in figures 5C,D,E. Here, ‘Labels’ is the lone optional name-value argument.

~~~
isomatrix_velocity(A);
isomatrix_velocity(A,1);
isomatrix_velocity(A,2);
isomatrix_velocity(A,3);
~~~

**Figure 4.**
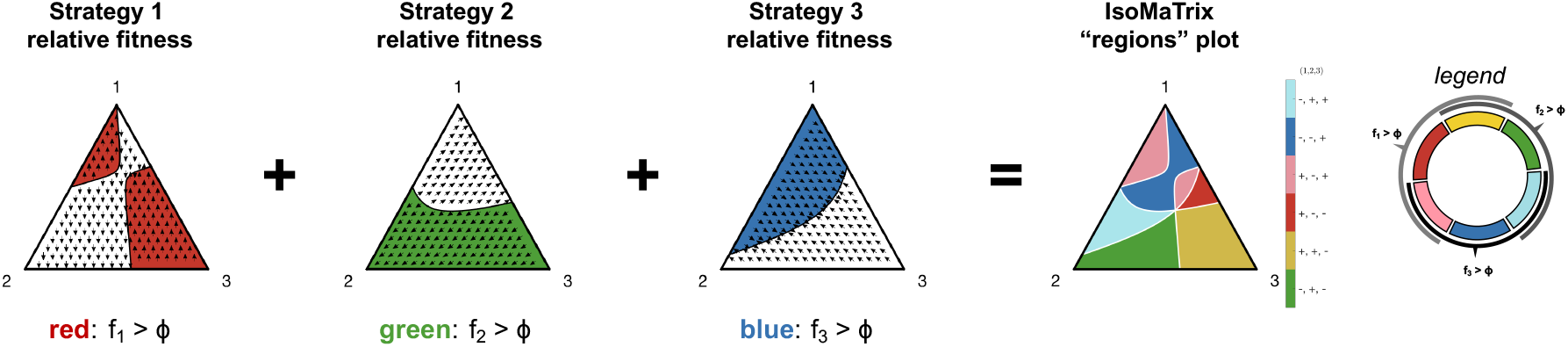
Schematic explanation for “isomatrix_region”. The region of the state space for which each strategy has positive relative fitness (*f*_*i*_ > *ϕ*) is colored in red (*f*_1_), green (*f*_2_), blue (*f*_3_). A “regions” plot is created by combining these into a single plot (right), where overlapping regions are colored as mixing colors (see legend).

**Figure 5.**
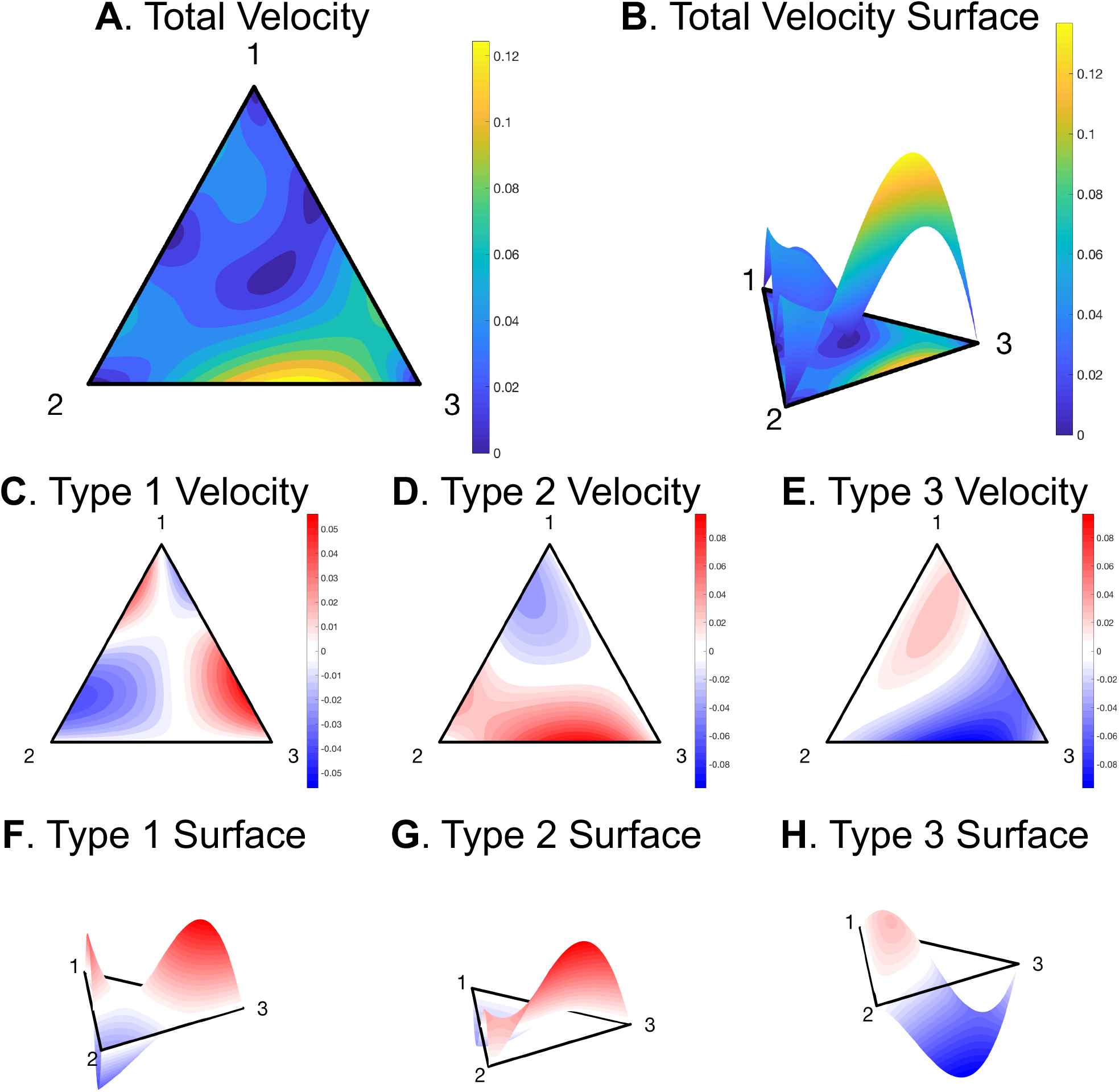
IsoMaTrix Velocity and Surface plot functions. (A) The output of “isomatrix_velocity” is shown for total velocity, which can also be displayed as a 3-dimensional surface plot, (B). (C,D,E) The velocity plots for each subtype can also be displayed as surface plots, shown in (F,G,H).

##### 1.7 isomatrix_fitness(A,id)

The purpose of this function is to color-code the background of the IsoMaTrix diagram according to the strategy fitness. The function takes as arguments a 3×3 payoff matrix *A*, and a strategy id. The id should be an integer (i.e. 1, 2, or 3) to specify which strategy fitness to compute, *f*_*i*_ = ∑ _*j*_ *a*_*i j*_*x* _*j*_. If no id is specified, the average fitness is computed: *ϕ* = ∑ _*j*_ *f* _*j*_*x* _*j*_. Here, ‘Labels’ is the lone optional name-value argument.

~~~
isomatrix_fitness(A);
isomatrix_fitness(A,1);
isomatrix_fitness(A,2);
isomatrix_fitness(A,3);
~~~

##### 1.8 isomatrix_region(A)

The region function divides the IsoMaTrix diagram into regions of positive or negative relative fitness (*f*_*i*_ − *ϕ*) of each strategy (delineated by the strategy isoclines in white). The signs indicated by colorbar. For example, the blue region in figure 2B indicates the region where *f*_1_ < *ϕ, f*_2_ < *ϕ*, and *f*_3_ > *ϕ* : (-, -, +). This diagram is also generated automatically using the “isomatrix” function.

~~~
isomatrix_region(A);
~~~

Other optional name-value arguments are ‘Labels,’ as well as ‘Color,’ ‘LineWidth,’ and ‘LineStyle’ used to specify the isoclines which bound the delineated regions. A schematic explanation which explains how an “isomatrix_region” is conceptually understood is seen in figure 4.

##### 1.9 isomatrix_surface(A,id)

This function extends “isomatrix_velocity” diagrams to a 3-dimensional surface plot. The arguments are identical, taking as arguments a 3×3 payoff matrix *A*, and a strategy “id” to specify which subtype velocity to compute. Examples for the following code are shown in figure 5.

~~~
isomatrix_surface(A);
isomatrix_surface(A,1);
isomatrix_surface(A,2);
isomatrix_surface(A,3);
~~~

##### 1.10 isomatrix_separatrix(A)

This function draws all internal separatrices associated with the replicator dynamics, and colors the background according to unique basins of attraction. There are at most three distinct basins of attraction possible in linear matrix games, resulting in at most three colors. The function returns a vector representing the fractional area each basin covers:

~~~
[red_area,green_area,blue_area]=isomatrix_separatrix(A);
~~~

Other optional name-value arguments are ‘Labels,’ as well as ‘Color,’ ‘LineWidth,’ and ‘LineStyle’ used to specify the style of each separatrix.

#### 2 IsoMaTrix Helper Functions (MATLAB)

##### 2.1 Coordinate transformations

Two functions are used internally for coordinate transformations. “XY_to_UVW(p)” takes in a vector or array of Cartesian coordinates (dimension: 2 by *n*) and converts to ternary coordinates. “UVW_to_XY(x)” takes in a vector or array of ternary coordinates (dimension: 3 by *n*) and converts to Cartesian coordinates for plotting.

##### 2.2 replicator(t,x,A)

The purpose of this function is to describe the coupled ordinary differential equations (eq. 1). This function is used for solving trajectories using MATLAB’s ode45 function.

##### 2.3 line_plot(A,x0,tF)

The purpose of this function is plot the matrix game’s trajectory over time on a line plot (x-axis is time, and y-axis is each strategy’s fraction over time). Here, ‘Labels’ is an optional argument used for legend entries. The following code was used to produce figure 6B:

~~~
tF=100;
x0=[0.3,0.3,0.4];
isomatrix_trajectory(A,x0,tF,Labels’,{‘1’,’2’,’3’});
~~~

**Figure 6.**
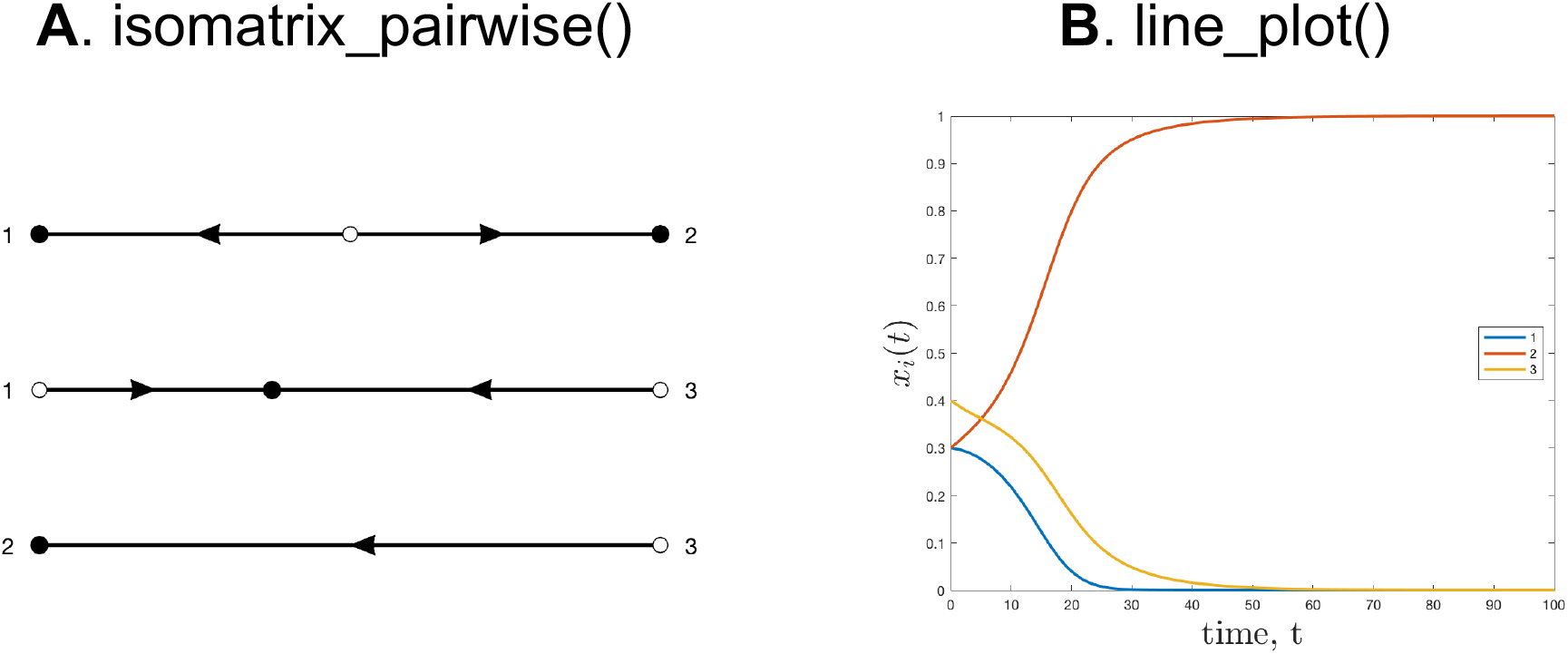
IsoMaTrix pairwise and line plot functions. (A) isomatrix_pairwise iterates through each pairwise two-strategy interaction and plots fixed points. (B) line_plot plots a trajectory over time.

##### 2.4 add_labels(string_array)

The purpose of this function is to add labels to the corners of the IsoMaTrix diagram, indicating the name of each type. It is best practice that these names are a single character or number. Alternatively labels can be specified as a name-value argument to most of the other isomatrix functions. This function takes as an argument a cellarray of 3 elements:

~~~
labels={‘1’,’2’,’3’};
add_labels(labels)
~~~

##### 2.5 add_gridlines(gridlines)

This function add gridlines to IsoMaTrix diagrams (step size of 1/gridlines), of a specified color. An example is shown in figure 8B.

~~~
add_gridlines(20);
~~~

##### 2.6 pairwise_fixedpoint(A)

In the case that a user desires to visualize the pairwise fixed points for a payoff matrix of arbitrary size, this function iterates and displays each pairwise interaction diagram. An example is shown in figure 6A. Optional name-value arguments are ‘Labels’, and ‘Color.’

~~~
isomatrix_pairwise(A);
~~~

##### 2.7 hessian(x,A)

This function calculates the Hessian matrix for a given fixed point, and is used internally for calculating stability of fixed points and determining basins of attraction.

##### 2.8 A_subset(A,types)

The purpose of this function is to plot a three-strategy subset of a larger payoff matrix. Given an *n* by *n* payoff matrix (*n* > 3), A_subset will output a 3 by 3 payoff matrix representing the competition values between the types specified in ‘types’ vector. Consider the following payoff matrix:

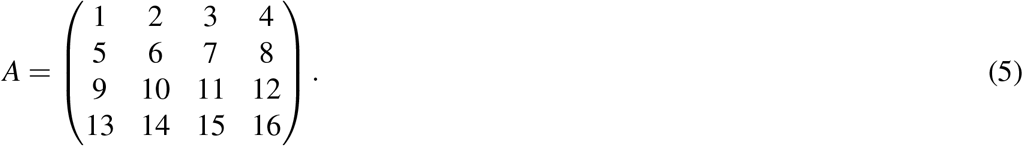

For example, consider the following lines of code:

~~~
B=A_subset(A,[1,2,4]);
~~~

This will produce the following output:

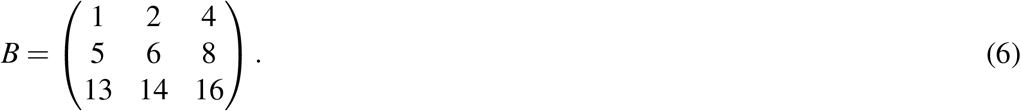

##### 2.9 Ohtsuki_Nowak_transform(A,k,rule)

This function transforms an *n*-by-*n* dimensional payoff matrix using the Ohtsuki-Nowak transformation. The transformation requires specifying the degree of graph, *k* (subject to the constraint that *k* > 3), and the update rule: “DB”(death-birth), “IM”: (immitation), or “BD” (birth-death)^29^. The returned matrix can then be passed to other IsoMaTrix functions for visualization.

#### 3 HAL integration with IsoMaTrix (Java)

The spatial games in this software package are simulated using the “Hybrid Automata Library” (HAL), a Java library developed for use in mathematical oncology modeling^26^. IsoMaTrix is bundled with version 1.1.0 of HAL. For installation instructions (and helpful hints on setting up an IDE for ease of computation) please visit HAL’s website for more information: http://halloworld.org/.

##### 3.1 Setting up Integrated Development Environment

Brief instructions are enclosed below for setting up a recommended Integrated Development Environment (IDE) for installing and running HALMatrixGames.

1. Download IsoMaTrix
2. Open Intellij Idea (a) click “Import Project” from the welcome window. (If the main editor window opens, Navigate to the File menu and click New -> “Project from Existing Sources”) (b) Navigate to the directory with the unzipped source code (“IsoMaTrix”). Click “Open.”
3. Intellij will now ask a series of questions/prompts. The first prompt will be “Import Project,” and you will select the bubble that indicates “Create project from existing sources” and then click “Next.”
4. The next prompt is to indicate which directory contains the existing sources. Navigate to the IsoMaTrix folder and leave the project name as “IsoMaTrix.” Click Next.
5. Intellij may alert you that it has found several source files automatically. Leave the box checked and click Next.
6. Intellij should have imported two Libraries: 1) lib and 2) HalColorSchemes. If these are not found, you’ll need complete the optional step 10 after setup is complete.
7. Intellij will prompt you to review the suggested module structure. This should state the path to the “IsoMaTrix” directory. Click next.
8. Intellij will ask you to select the Java JDK. Click the “+” and add the following files: (a) Mac: navigate to “/Library/ Java/ JavaVirtualMachines/” (b) Windows: navigate to “C: Program Files Java” (c) Choose a JDK version 1.8 or later
9. Intellij will state “No frameworks detected.” Click Finish.
10. If step 6 failed, you will need to do one more step and add libraries for 2D and 3D OpenGL visualization. Navigate to the File menu and click “Project Structure.” Click the “Libraries” tab. Use the minus button (-) to remove any pre-existing library entries. Click the “+” button, then click “Java” and direct the file browser to the “IsoMaTrix/HAL/lib” folder. Click apply or OK.

##### 3.2 HALMatrixGame2D and HALMatrixGame3D

Two nearly identical classes are supplied in the HALMatrixGame java code. HALMatrixGame2D simulates matrix games on two-dimensional lattice grids, and HALMatrixGame3D extends the domain to three-dimensions. To simulate a matrix game, run the main function of either class.

These two classes on are built in Hybrid Automata Library (HAL)^26^, as extensions of AgentGrid2D and AgentGrid3D, respectively. Each lattice point within the grid contains exactly one agent, called Cell2D or Cell3D (extension of HAL’s AgentSQ2D and AgentSQ3D classes, respectively).

Both of the HALMatrixGame classes utilize the “imitation updating” rule^33^. At each time step a randomly chosen cell (the focal cell) updates its strategy to imitate one of its own neighbors in proportion to fitness. This update rule include the focal cell within the calculation, so it may ‘imitate’ its own strategy if it is the most fit. The following specifications are available to the user:

##### 3.3 Fitness Neighborhood

The neighborhood of cells used to calculate the fitness of the focal cell may also be specified. For example, HAL includes von Neumann neighborhood (nearest 4 neighbors up, left, down, right) or Moore neighborhood (nearest 8 neighbors which include the von Neumann cells with diagonals included). These are specified in the following way:

~~~
int[]neighborhood=VonNeumannHood(true);
~~~

or,

~~~
int[]neighborhood=MooreHood(true);
~~~

Note: the boolean argument for each is set to true, indicating consideration of the focal cell in the neighorhood. This is required for imitation updating. Examples are shown in figure 7.

**Figure 7.**
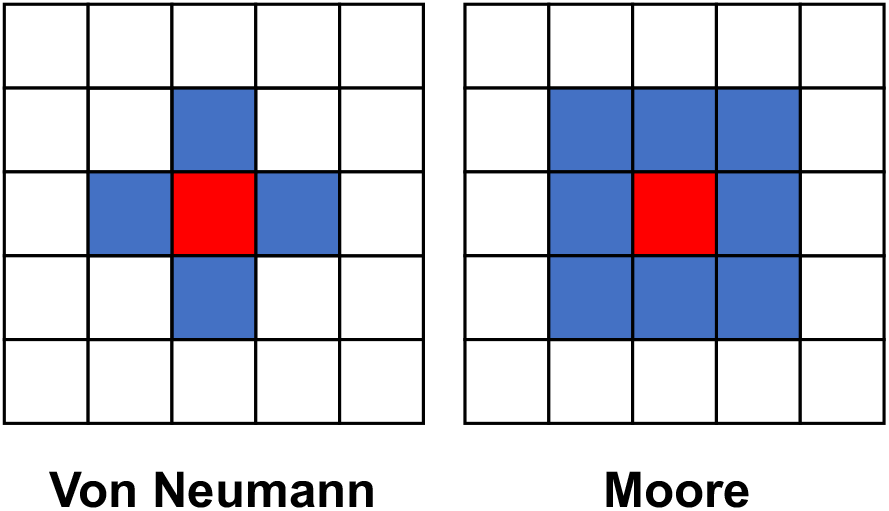
HalMatrixGames neighborhoods. Fitness is calculated by interactions within the neighborhood (blue) of the focal cell (red). Two options are a Von Neuman (4 neighbors) and a Moore (8 neighbors).

##### 3.4 Deterministic or Stochastic Updating

Deterministic updating selects the most fit individual within the neighborhood and updates the strategy of the focal cell to match the strategy of the most fit individual. In case of tie, a randomly selected individual is chosen between those that tie. Stochastic updating updates according to a probability density function weighted by the fitness of each neighbor. This option is chosen by specifying the “PROCESS” parameter in java:

~~~
public int PROCESS = DETERMINISTIC;
~~~

or,

~~~
public int PROCESS = STOCHASTIC;
~~~

##### 3.5 Population Update Fraction

Each time step, a fraction of the population is selected to undergo the imitation update replacement process. This parameter is called ‘UPDATE_FRACTION’ and is bounded between 0 and 1 (inclusive).

~~~
public double UPDATE_FRACTION = 1.0;
~~~

##### 3.6 SingleSimulation(int timesteps)

The purpose of this function is to simulate the dynamics of a single matrix game on a two- or three-dimension grid with a specified initial fraction of each substrategy, 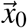. The dynamics are simulated for specified number of timesteps, and can be easily visualized on an IsoMatrix diagram using the “HAL_isomatrix_trajectory” function (MATLAB).

Note: the data from the simulation in SingleSimulation is saved in the “HALMatrix-output” folder, in a file called “HAL_trajectory.csv,” which is automatically utilized when the HAL_isomatrix_trajectory function is called in MATLAB. If this file is renamed by the user, it must be re-specified in HAL_isomatrix_trajectory. An example of this function is shown in “StartHere.java”:

~~~
int sideLength = 20;
HALMatrixGame2D matrixGame = new HALMatrixGame2D(sideLength);

// fraction of population
matrixGame.UPDATE_FRACTION = 0.5;
matrixGame.PROCESS = DETERMINISTIC;
matrixGame.payoffs = new double[]{
       0.7,0.0,0.7,
       0.3,0.4,0.8,
       1.0,0.3,0.2};

int timesteps = 100;
matrixGame.SingleSimulation(timesteps);
~~~

##### 3.7 MeshGrid(int timesteps, int nSims)

The purpose of this function is to simulate a “meshgrid” of single matrix game simulations which can later be combined to produce quiver, velocity, uncertainty, and region plots in IsoMaTrix, in MATLAB. The function takes as arguments a side length (integer), number of time steps (integer), and number of stochastic simulations per grid point.

The side length is an integer specifying the domain size in 2- or 3-dimensions. Dynamics are simulated for an evenly distribution of initial conditions (figure 8B). The average velocity will calculated by subtracting the final state vector, 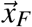 from the initial state vector, 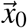, shown in figure 8C, and described in more detail in the “HAL_isomatrix_quiver” section.

**Figure 8.**
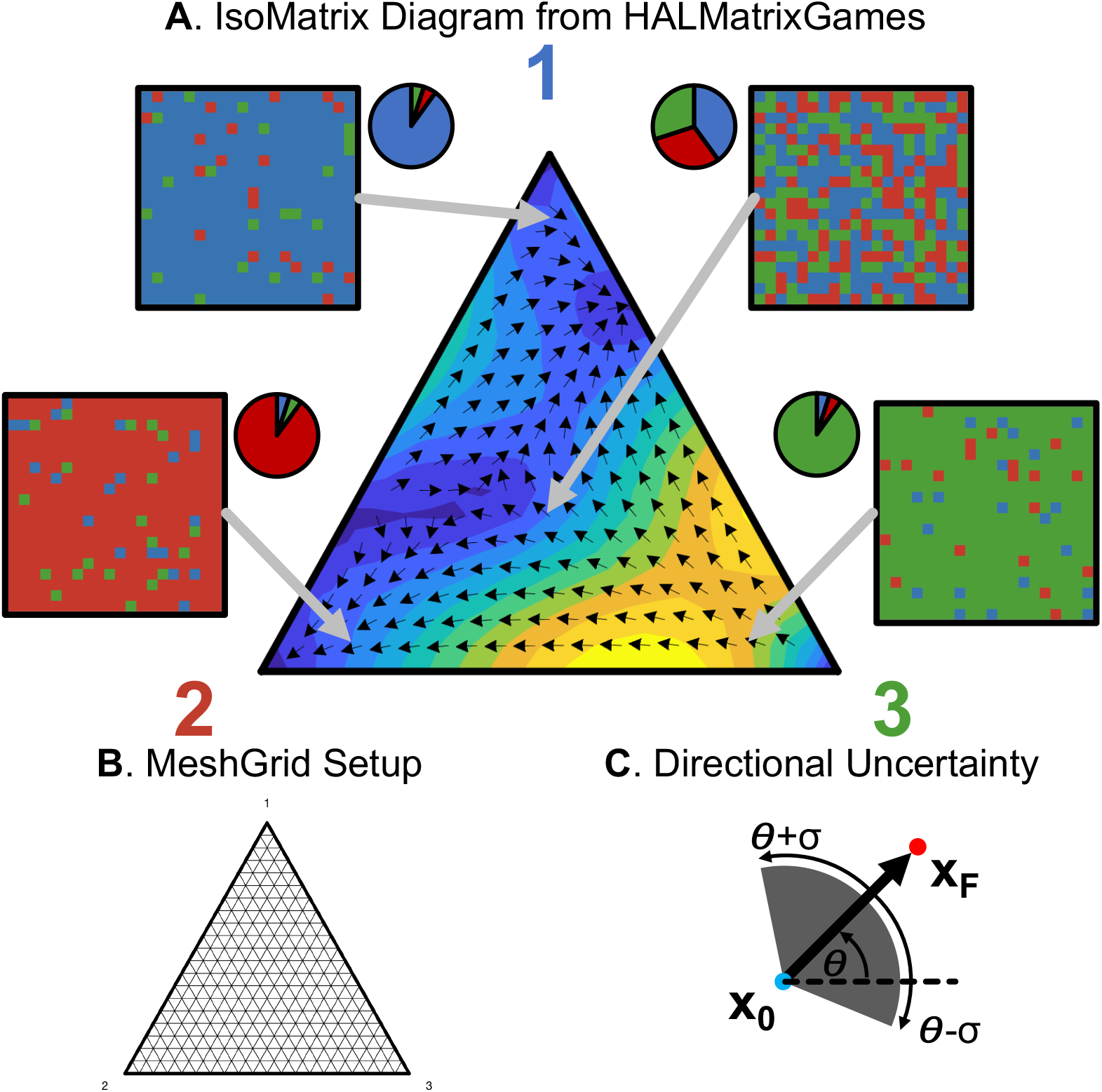
HalMatrixGames in IsoMaTrix. (A) Schematic of HALMatrixGames visualized on an IsoMaTrix diagram. (B) MeshGrid Setup. (C) Directional uncertainty of resultant 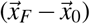 vector, R. The gray arc shows one standard deviation in each direction: *±σ* (see eqn. 8.)

Note: the data from the simulations in MeshGrid is saved in the “HALMatrix-output” folder, in a file called “IsoMaTrixGrid.csv,” which is automatically read into the relevant isomatrix functions described in the next section (HAL_isomatrix_quiver, HAL_isomatrix_region, HAL_isomatrix_trajectory, HAL_isomatrix_velocity, and HAL_isomatrix_uncertainty). If this file is renamed by the user, it must be re-specified in each of these functions. An example of this function is shown in “StartHere.java”:

~~~
int side_length = 20;
int time_steps = 1;
int nSims = 50;
HALMatrixGame2D model = new HALMatrixGame2D(side_length);
model.MeshGrid(time_steps,nSims);
~~~

#### 4 Visualizing HALMatrixGames using IsoMaTrix

As mentioned in the previous section, HALMatrixGames generate CSV files interpretable by IsoMaTrix functions (in MATLAB) to generate corresponding IsoMaTrix diagrams. The following sections describe the functions used to display quiver, region, velocity, or uncertainty plots *after* simulations are performed in HALMatrixGames. A schematic of the setup is shown in figure 8. HALMatrixGames are initialized for each initial proportion (8A, inset two-dimensional lattices), across a meshgrid of initial conditions (8B). Velocity can be estimated by the resultant vector which subtracts the initial condition, 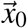, from the final proportion, 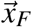, after user-specified number of time steps (8C).

##### 4.1 HAL_isomatrix()

HAL_isomatrix is the canonical function to display results of spatial evolutionary dynamics on IsoMaTrix diagrams. Example output is shown in figure 9 (with deterministic updating, update fraction of 1).

**Figure 9.**
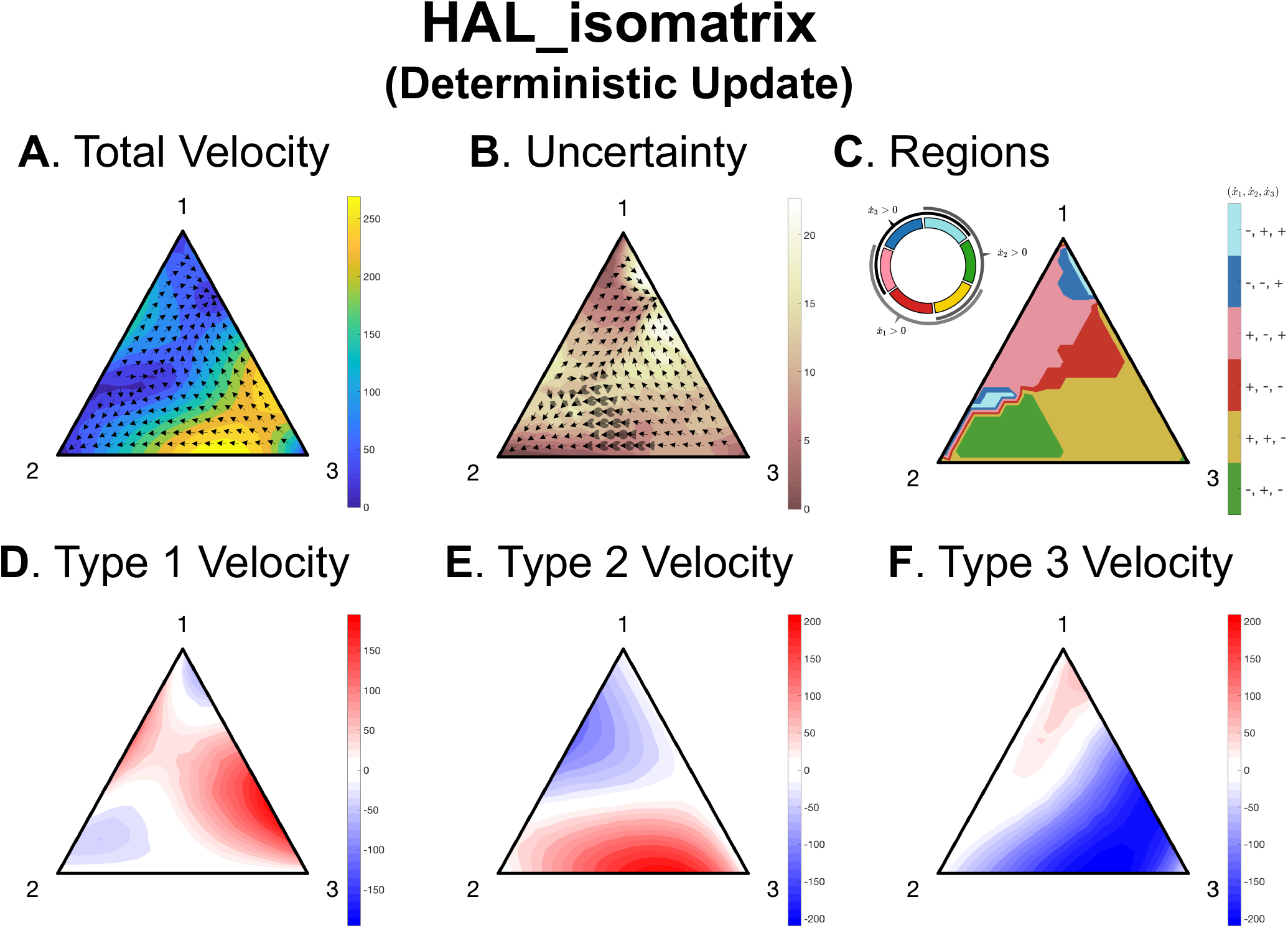
HalMatrixGames in IsoMaTrix with the deterministic update rule. (A) A quiver plot (black arrows) shows the phase flow, where the background color indicates the magnitude of the velocity vector. (B) The same quiver plot, with uncertainty in velocity direction shown by arcs shown by transparent arcs on each arrow, and uncertainty in velocity magnitude shown by background color. (C) A “region” plot divides the diagram into regions of positive or negative growth of each strategy, with the signs indicated by colorbar. For example green indicates the region where 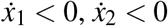, and 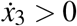. (C,D,E) Strategy velocity magnitude is color-coded by positive (red) or negative (blue) growth of each strategy, with nullcline shown in black.

HAL_isomatrix();

This automatically generates a series of figures, corresponding to figure 9A-F:

**A** quiver & total velocity

**B** quiver & total velocity uncertainty

**C** “regions” plot

**D** subvelocity for strategy 1

**E** subvelocity for strategy 2

**F** subvelocity for strategy 3

This makes for easy comparison between spatial evolutionary dynamics (figure 9) and well-mixed, replicator dynamics (figure 2). Optionally, the user may also specify the isomatrix ‘Labels’ and ‘Filename’ (filename is required if “MESH_GRID_FILENAME” is changed in StartHere.java).

~~~
HAL_isomatrix(‘Filename’,’IsomatrixGrid.csv’, … ‘Labels’,{‘1’,’2’,’3’});
~~~

##### 4.2 HAL_isomatrix_trajectory(color)

This MATLAB function is used to plot the “SingleSimulation” function output onto an IsoMaTrix diagram. Optionally, a filepath can be specified to if the “SINGLE_SIMULATION_FILENAME” is altered from the default (HAL_trajectory.csv).

~~~
HAL_isomatrix_trajectory();
~~~

or, specifying the optional name-value arguments as follows:

~~~
HAL_isomatrix_trajectory(‘Color’,red,’Labels’,{‘1’,’2’,’3’}, …
               ‘LineWidth’, 2,’LineStyle’, ‘:’, …
               ‘Filename’, ‘HAL_trajectory.csv’);
~~~

If the file was previously renamed by the user, a new filepath must be specified. If no color is specified, the trajectory will be black, solid, with linewidth of 2.

##### 4.3 HAL_isomatrix_quiver(uncertainty_boolean)

This MATLAB function is used to plot the “MeshGrid” function output onto an IsoMaTrix diagram. As described in the MeshGrid section, simulations are initialized from an evenly distributed grid (figure 8B). The velocity is estimated by subtracting the initial fraction from the final fraction, 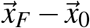 for each stochastic simulation in HALMatrixGame (see figure 8C). The function takes an optional argument of “uncertainty_boolean” which indicates if the uncertainty cone should be drawn about each quiver arrow, shown in figure 8C (default value of false).

~~~
HAL_isomatrix_quiver(false);
~~~

Note: this MATLAB function reads in the “MESH_GRID_FILENAME” (default name is “IsoMaTrixGrid.csv”) file from the “MeshGrid” of simulations in HALMatrixGames, and the filepath must be specified if this file is renamed or moved. If no color is specified, the quiver arrows will be black.

~~~
HAL_isomatrix_quiver(false,’Color’,black, …
                      ‘Filename’, ‘IsoMaTrixGrid.csv’, …
                      ‘Labels’,{‘1’,’2’,’3’});
~~~

##### 4.4 HAL_isomatrix_velocity(id)

The purpose of this function is to color-code the background of the IsoMaTrix diagram according to the average velocity of the HALMatrixGames simulations. The function takes an argument a strategy id (i.e. 1, 2, or 3) to specify which strategy velocity to compute. The resultant velocity vector is estimated by subtracting the initial fraction from the final fraction, 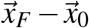 for each stochastic simulation in HALMatrixGame (figure 8C). If no strategy id is specified, the magnitude of the resultant velocity, ‖ *v* ‖, of all strategies is computed. Let 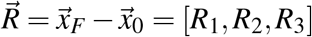.

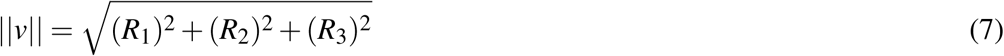

The following lines of code can be used to produce the background color of figure 9A,C,D, and E for total velocity, strategy 1, 2, and 3 respectively:

~~~
HAL_isomatrix_velocity();
HAL_isomatrix_velocity(1);
HAL_isomatrix_velocity(2);
HAL_isomatrix_velocity(3);
~~~

Note: this MATLAB function reads in the “MESH_GRID_FILENAME” (default name is “IsoMaTrixGrid.csv”) file from the “MeshGrid” of simulations in HALMatrixGames, and the filepath must be specified if this file is renamed or moved.

~~~
HAL_isomatrix_velocity(‘Filename’, ‘IsoMaTrixGrid.csv’, …
                       ‘Labels’,{‘1’,’2’,’3’});
~~~

##### 4.5 HAL_isomatrix_region()

The purpose of this function is to divide the IsoMaTrix diagram into regions of positive or negative growth of each substrategy. The signs indicated by colorbar. For example, the green region in figure 9C indicates the region where 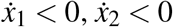, and 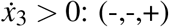. The velocity is estimated by subtracting the initial fraction from the final fraction, 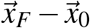 for each stochastic simulation in HALMatrixGame (see figure 8C).

~~~
HAL_isomatrix_region();
~~~

Note: this MATLAB function reads in the “MESH_GRID_FILENAME” (default name is “IsoMaTrixGrid.csv”) file from the “MeshGrid” of simulations in HALMatrixGames, and the filepath must be specified if this file is renamed or moved.

~~~
HAL_isomatrix_region(‘Filename’, ‘IsoMaTrixGrid.csv’, … ‘Labels’,{‘1’,’2’,’3’});
~~~

##### 4.6 HAL_isomatrix_uncertainty(id)

The purpose of this function is to color-code the background of an IsoMaTrix diagram corresponding to the standard deviation of the magnitude of the estimated velocity vector. The velocity is estimated by subtracting the initial fraction from the final fraction, 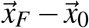 for each stochastic simulation in HALMatrixGame (see figure 8C. The function takes as an argument a strategy id. The id should be an integer (i.e. 1, 2, or 3) to specify which strategy velocity uncertainty to compute. If no strategy id is specified, the magnitude of the resultant velocity, ‖ *v* ‖, of all strategies is computed (eqn. 7).

~~~
HAL_isomatrix_uncertainty();
~~~

The uncertainty is the standard deviation of the magnitude of the velocity vector, calculated using MATLAB’s “std” function. This defines the standard deviation of vector, 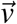, consisting of *i* = 1, …, *M* stochastic realizations as:

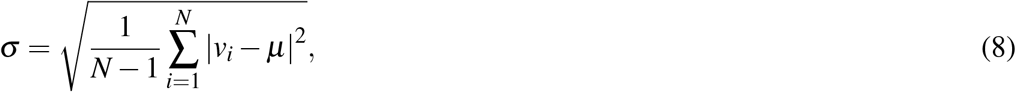

where *μ* is the mean:

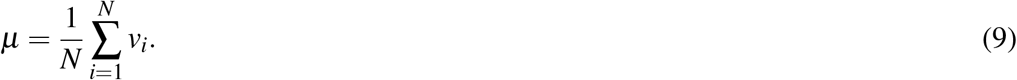

Note: “HAL_isomatrix_uncertainty” displays the uncertainty of the magnitude of the velocity vector (i.e. eqn. 7), while the uncertainty in the direction of the resultant vector is displayed in “HAL_isomatrix_quiver,” shown in figure 8C.

Note: this MATLAB function reads in the “MESH_GRID_FILENAME” (default name is “IsoMaTrixGrid.csv”) file from the “MeshGrid” of simulations in HALMatrixGames, and the filepath must be specified if this file is renamed or moved.

~~~
HAL_isomatrix_uncertainty(‘Filename’, ‘IsoMaTrixGrid.csv’, … ‘Labels’,{‘1’,’2’,’3’});
~~~

## Notes

https://github.com/MathOnco/isomatrix

### Competing Interest Statement

The authors have declared no competing interest.

